# Insertional mutagenesis defines drivers and evolutionary relationships in pancreatic cancer metastasis

**DOI:** 10.1101/2020.09.28.317701

**Authors:** Suman Govindaraju, Michelle Maurin, Justin Y. Newberg, Devin J. Jones, Hubert Lee, Judit E. Markovitz, Michael B. Mann, Michael A. Black, Karen M. Mann

## Abstract

Metastasis is a defining feature of pancreatic cancer, impacting patient quality of life and therapeutic outcomes. While recurrent mutations in *KRAS* and *TP53* play important roles in primary tumor development and persist in metastases, the contributions of other genes to the metastatic process is understudied. Here, we define a network of metastasis-promoting genes and uncover positively selected genes in metastatic lesions from our *Sleeping Beauty* mouse model of pancreatic cancer using our recently described SB Driver Analysis statistical pipeline. We show that loss of single genes *DLG1, PARD3, PKP4* and *PTPRK* promote pancreas cancer progression in a context-specific manner. Finally, we define clonal and sub-clonal insertion events that distinguish primary and individual metastatic tumors and use single cell sequencing to gain insight into the evolutionary relationships between sub-regions of primary pancreatic tumors and related metastases, which has important implications for pancreatic disease progression and therapeutic opportunities for patients.

## INTRODUCTION

Pancreatic cancer is a highly metastatic disease with the second-highest mortality rate behind lung cancer (Siegel, Miller, & Jemal, 2019). Because the majority of patients are diagnosed at late disease stages, there is an urgent need to better understand the cellular programs that promote pancreatic cancer metastases and to uncover molecular vulnerabilities of metastases that can be pursued therapeutically. Genomic and transcriptomic analyses of primary pancreatic cancers and in limited metastatic samples have demonstrated the paucity of recurrent mutations in pancreatic cancer outside of oncogenic *KRAS* activation and loss of function of *CDKN2A, TP53, SMAD4* and *BRCA1* (Biankin et al., 2012; Witkiewicz et al., 2015; Wood et al., 2007). In particular, *SMAD4* loss, occurring in approximately 50% of patients, is considered a latent event and is associated with an increased incidence of distal metastasis (Iacobuzio-Donahue et al., 2000). Recently, Connor *et al.* reported that *KRAS* amplification and increased mutant *TP53* expression are important features of liver metastases (Connor et al., 2019), supporting earlier in-depth genomic sequencing analysis of primary pancreatic tumors and related metastatic lesions (Yachida et al., 2010). Further, a widely-studied mouse model of pancreatic cancer driven by the cooperation of an oncogenic *Kras^G12D^* mutation and a gain-of-function *Trp53* mutation demonstrated that these two mutations are necessary and sufficient to drive pancreatic cancer in the mouse. However, additional genetic and epigenetic events are required to drive metastasis (Campbell et al., 2010) and little is understood about the timing and cooperation of genetic and epigenetic events to promote advanced PDAC.

Spontaneous mouse models of cancer are powerful tools to gain insight into the underlying genetic aberrations that initiate, promote and maintain metastatic disease. Two genetic screens using the *Sleeping Beauty* insertional mutagenesis system in the background of oncogenic *Kras^G12D^* uncovered hundreds of cooperating candidate drivers, many of them novel, including *Ctnnd1* and *Usp9x* (Mann et al., 2012; Perez-Mancera et al., 2012). However, the extensive list of candidate drivers confounded approaches to prioritize and validate these genes for their biological role in promoting disease. One hallmark of the SB pancreatic cancer (SB_PDAC) mouse model is that several mice developed aggressive, metastatic disease which appeared in part to be influenced by the SB transposon, as all mice with the T2/Onc3 transposon (Mann et al., 2012) developed metastasis, while only half of the mice driven by the T2/Onc2 transposon exhibited lesions outside the pancreas. Further, bulk sequencing of SB insertions in pancreatic tumors revealed a high degree of inter- and intra-tumor heterogeneity (ITH), modeling a salient feature of human pancreatic cancers. ITH has been postulated to influence disease development and therapeutic resistance in several cancers, including breast, colon, lung, head and neck, melanoma and pancreatic (Hua et al., 2020; Jamal-Hanjani, Quezada, Larkin, & Swanton, 2015; McDonald et al., 2019; Mroz et al., 2013; Niknafs et al., 2019; Reuben et al., 2017; Sveen et al., 2016; Zhao, Hemann, & Lauffenburger, 2014).

Recently, we developed a statistical analysis pipeline, termed SB Driver Analysis (Newberg, Black, et al., 2018), to define driver genes in sequencing datasets from *Sleeping Beauty* mouse models of human cancers, enabling the evaluation of cohorts driven by independent transposons (Newberg, Mann, Mann, Jenkins, & Copeland, 2018) and applicable across data generated from several NGS platforms, including Roche 454, Illumina HiSeq and ThermoFisher Ion Torrent (Mann et al., 2016). Here, we applied the SB Driver Analysis pipeline to define early-selected drivers in pancreatic cancer, define drivers of metastasis, explore the impact of intra-tumor heterogeneity on disease progress and perform *in vitro* and *in vivo* functional validation experiments to demonstrate the biological impact of novel genetic drivers in promoting metastatic disease.

## RESULTS

### Identification of Early Progression Drivers in pancreatic cancer

Pancreatic cancers characteristically demonstrate a low mutation burden and few recurrent mutations outside of activating missense mutations in *KRAS* and inactivating mutations in latent drivers *TP53, CDKN2A* and *SMAD4.* We and others (Mann et al., 2012; Perez-Mancera et al., 2012) previously performed forward genetic screens using *Sleeping Beauty* insertional mutagenesis to identify additional loci that cooperate with the oncogenic *Kras*^G12D^ mutation to promote disease development in a mouse model of pancreatic cancer. These two studies utilized the same inducible *Kras*^G12D^ allele (Jackson et al., 2001) but independent SB transposons and different alleles of both the SB transposase and *Pdx1-cre.* Recently, we reanalyzed these insertion datasets, performing a combined analysis of the primary tumor data from each study using the gene centric SB Driver Analysis pipeline we recently developed (Newberg, Black, et al., 2018) to enhance statistical power to detect highly recurrent drivers (based on frequency of insertions) and novel drivers not found in each cohort independently. The 814 genes identified by combined driver analysis were reported in the SB Cancer Driver Database (SBCDDB (Newberg, Mann, et al., 2018)). The drivers from the combined analysis overlapped with two-thirds of the progression drivers from the independent cohort analyses reanalyzed with the SB driver analysis. Only twenty-nine drivers were defined from the combined analysis that were not found in either independent dataset analysis. These results suggested that the major drivers of the SB_PDAC model were captured in each independent study, while the combined analysis further refined the recurrent driver list for biological relevance to PDAC development.

Because we specifically want to harness these datasets to define recurrent, novel drivers that contribute to pancreatic cancer progression to adenocarcinoma and metastasis, here we have further refined our analysis to filter away insertions with low read depths prior to SB Driver Analysis on the individual and combined datasets, reasoning these are rare or background (unselected) events in the individual tumors. The combined analysis identified 115 recurrently altered drivers with high read depths (**Supplemental Table 1**). We termed these genes early progression drivers (EPDs), as they are found to occur with high frequency in several tumors within a cohort and are likely to be clonally selected during disease progression. For some drivers, there was a wide range of insertion read depths within the cohort, whereby a subset of tumors had insertions with very high read depths (greater than 100), indicative of highly clonal events, while other tumors with insertions in the same loci were represented by fewer than 5 reads. This suggests that clonal selection of these loci is highly heterogeneous across tumors. An oncoprint illustrating the heterogeneity of EPDs across the tumor cohort is shown in **Figure 1, panel A** (10 tumors or greater) and **Supplemental Figure 1**. The majority of the EPDs are predicted tumor suppressors based on the presence of insertions in both the sense direction (blue bars) and antisense direction (red bars) along the coding region. Seventy-eight EPDs were previously identified as progression drivers in our combined primary SB driver analysis at a frequency of 15% or greater, suggesting these drivers are positively and recurrently selected in SB_PDAC tumors in a clonal or sub-clonal manner. Further, the EPD analysis identified 21 new drivers. Thus, by filtering the sequence data for high read-depth insertions prior to driver analysis, we were able to uncover recurrent and highly statistically significant drivers of pancreatic cancer progression.

**Figure 1.**
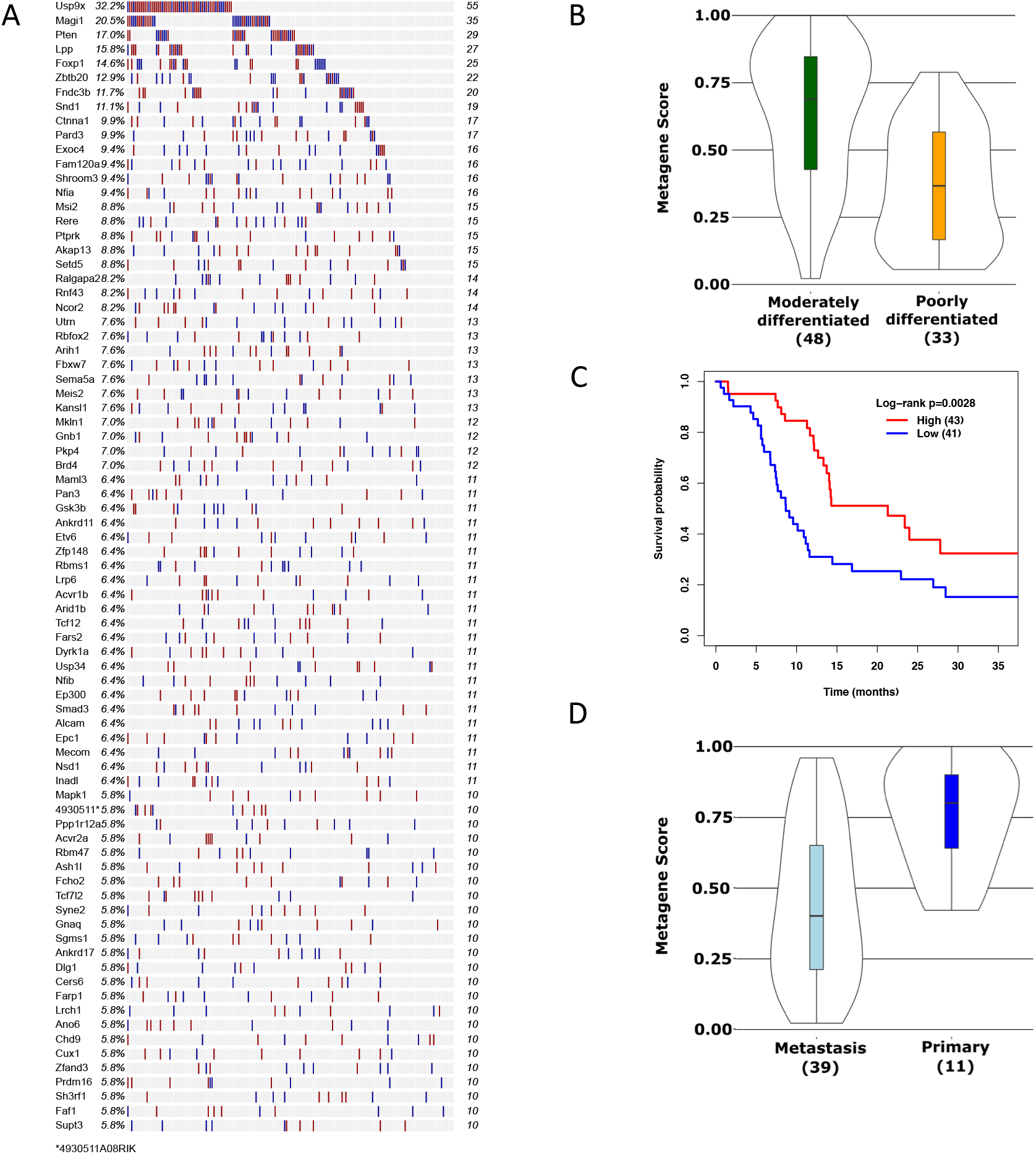
Early progression drivers in SB_PDAC tumors. An oncoprint of the top 79 early progression drivers (EPDs) (A) shows the incidence of SB insertions occurring in greater than 10 SB_PDAC tumors across the cohort of 172 primary tumors. Individual tumors (animals) are represented by the columns. Insertions present in the sense (blue) or antisense (red) strand depicted by the bars. (B) An expression metagene constructed from the human orthologs of SB_PDAC EPDs with RNASeq expression data (ICGCV28) is significantly associated with poorly differentiated primary tumor grade compared to moderately differentiated (t-test, *P*<0.001). (C) Low expression of the EPD metagene (67% quantile) in primary resected tumors is significantly associated with decreased disease-free survival (log-rank *P*-value=0.0028). (D) Expression of the EPD metagene is significantly decreased in pancreatic cancer liver metastases compared to primary tumors (ICGC EGA 3584, *P*=0.008).

### EPDs are implicated in human pancreatic cancers

We next interrogated human PDAC sequencing datasets (Bailey et al., 2016; Network, 2017; Witkiewicz et al., 2015) obtained from cBioPortal (Cerami et al., 2012) for the presence of mutations in 108 human orthologs of the SB_PDAC EPDs and observed that nearly all the EPDs had a least one mutation identified in a human PDAC genome, while 20 EPDs were mutated at a frequency greater than 1% (see oncoprint, **Supplemental Figure 2**). Missense mutations were the most common mutation type, followed by truncation. *RNF43* had the highest mutation frequency of 5%. *RNF43* encodes a negative regulator of Wnt signaling and mutations in this gene have been identified in human colon cancer patients (Eto et al., 2018; Neumeyer et al., 2019; Yaeger et al., 2018) and more recently associated with intraductal pancreatic mucinous neoplasms (IPMNs) of the pancreas (Furukawa et al., 2011; Sakamoto et al., 2015) and in PDAC genomes (Network, 2017). *Rnf43* was a driver also identified in an SB model of intestinal cancer (Takeda et al., 2016) suggesting that SB inactivation of this gene promotes intestinal and pancreatic cancers in the mouse. The low mutation frequency in the EPDs reflects the overall low recurrence of mutations in pancreatic cancer genomes, revealed through the sequencing efforts of resected primary tumors from several groups (Biankin et al., 2012; Connor et al., 2019; Network, 2017; Waddell et al., 2015). Because SB insertions disrupt gene expression, we predicted that these EPDs would be differentially expressed in human PDAC. RNASeq data from primary resected pancreas tumors from ICGC identified differential expression of EPDs across a subset of human PDAC tumors (see heatmap, **Supplemental Figure 3**). Importantly, we found that an expression metagene derived from these tumors showed significantly decreased expression of the EPDs associated with poorly differentiated tumors (**Figure 1, panel B,** t-test, *P*<0.001), which are more aggressive than well-differentiated tumors (Bailey et al., 2016; Collisson et al., 2011; Moffitt et al., 2015). Further, we observed a significant decrease in disease-free survival when comparing patient outcomes with EPD expression profiles representing the lower two-thirds expressing tumors to the outcomes for patients with the top one-third higher expressing tumors (log-rank *P*=0.0028, **Figure 1 panel C**). The Cox proportional hazards (CPH) *P*-values for all EPDs in the expression metagene are shown in **Supplemental Table 2**. We next investigated expression of the EPDs in liver metastases compared to primary tumors using RNASeq data from 35 liver pancreatic cancer metastases and 11 unmatched primary pancreatic tumors from the COMPASS trial (EGAD0000003584). The samples were equally matched for gender. We constructed a metagene for EPD expression (**Supplemental Figure 4**) and found a significant decrease in the EPD metagene expression in liver metastases compared to primary tumors (Rank-based metagene, F statistic *P*=0.008, **Figure 1 panel D**). Combined, these data suggest that reduced expression of EPDs plays a biological role in promoting PDAC liver metastases.

### Prioritization of EPDs for functional validation

Ninety-four of the EPDs mapped to at least one signaling pathway (Enrichr gene lists and Gene Ontology terms) with significant enrichment of drivers by GOSeq analysis in histone methyltransferase activity (*P*=3.5E-04), protein acetylation (*P*=3.3E-04), adherens junctions (*P*=6.7E-04) and WNT (R-HSA-4791275, *P*=0.03, **Supplemental Table 3**). Comparative genomic analyses have shown that despite molecular heterogeneity across tumors, mutated drivers converge on the same curated signaling pathways, suggesting that the genes *per se* are less important than the pathways they regulate. Further, identifying signaling nodes with potential intervention points for therapeutic opportunities can be enhanced by considering genetic interaction networks. Indeed, the SB_PDAC EPDs show a strong interaction network with few nodes connecting the majority of the drivers. In addition, the proteins encoded by 90 of the EPDs demonstrated more interactions than expected by chance (STRING, *P*<1.0E-16)(Szklarczyk et al., 2019), with major nodes at PTEN, EP300 and SMAD3 and secondary nodes MAPK1 (ERK2), SETD2 and PARD3 (**Supplemental Figure 5**). This PPI network highlights an enrichment of proteins involved in transcriptional regulation, including transcription factors and chromatin modifying enzymes, in addition to several kinases, ubiquitinases and scaffolding proteins and in members of the Hippo pathway (KEGG, *P*_FDR-corrected_=7.35E-06, STRING). Together, these data demonstrate that the EPDs defined by the SB_PDAC models have biological interactions in important signaling pathways and processes relevant to human PDAC.

We prioritized 60 EPDs for gene expression analysis using a combination of Nanostring (**Supplemental Figure 6**) and qPCR assays (**Supplemental Figure 7 and Supplemental Table 4**), across 11 human PDAC cell lines with epithelial or mesenchymal properties derived from primary tumors, metastatic sites or from immortalized and transformed pancreas ductal cells. We included 16 additional genes that were defined progression drivers private to PDAC or were frequent in SB_PDAC models as a control gene set. Using hierarchical clustering of the Nanostring data, we observed distinct expression patterns across the cell lines (**Supplemental Figure 6, panel A**). Cell lines PATC148 and PATC153 were derived from PDX models of primary tumors (Kang et al., 2015) from individual patients with liver metastases. PATC148 contains a *KRAS*^G12D^ mutation while PATC153 is *KRAS* wild type; PATC43 was derived from a PDX mouse model of a PDAC tumor with a *KRAS*^G12V^ mutation (J. Fleming, personal communication). Expression of epithelial marker E-cadherin and mesenchymal marker vimentin defined the epithelial and mesenchymal properties of the cell lines (**Supplemental Figure 6, panels B and C**). Cell lines expressing both markers are defined as quasi-mesenchymal or basal (Bailey et al., 2016; Collisson et al., 2011; Moffitt et al., 2015). Further, principle component analysis illustrated distinct expression patterns of EPDs between epithelial and mesenchymal cell lines (**Supplemental Figure 6, panel D**). *PTPRK* and *FOXP1* are more highly expressed in epithelial-like cell lines, while *RBFOX2* is more highly expressed in mesenchymal-like cell lines. Expression of genes involved in Hippo signaling was varied. While *PARD3* showed relatively high expression in PATC153 and AsPC1 cells, *INADL* (*PATJ*) was highly expressed in epithelial-like cells and lower in mesenchymal-like cells; *DLG1* and *TAOK1* expression measured by qPCR was downregulated in most cell lines compared to HPNE control cells (**Supplemental Figure 7, panels D and G**).

### In vitro and in vivo validation of EPDs in PDAC progression

Utilizing our focused PPI network and our cell line expression data, we chose 10 novel drivers for functional validation studies in two or more human PDAC cell lines, using two independent shRNAs to knockdown individual genes or pooled shRNAs against select single targets (**Supplemental Table 4**), and successfully achieved 2-fold or greater knockdown for 8 drivers. We chose the cell lines expressing the same shRNA that achieved the highest level of knockdown and further characterized 4 EPDs (*DLG1, PARD3, PKP4, PTPRK*) in epithelial-like and mesenchymal-like cells. We found that depletion of *DLG1* (**Figure 2, panel A**) increased cellular proliferation in epithelial-like PL45 cells, with no change in quasi-mesenchymal Panc1 cells (**Figure 2, panels B and C**). However, cell migration significantly decreased in PL45 cells (**Figure 2, panel D**) and significantly increased in Panc1 cells (**Figure 2, panel E**), illustrating the context specificity of *DLG1* loss in promoting aggressive phenotypes. *DLG1* encodes a scaffolding protein containing PDZ domains that is part of the Scribble polarity module (Bonello & Peifer, 2019), plays a role in the formation of adherens junctions in intestinal cells (Laprise, Viel, & Rivard, 2004) and negatively regulates cell proliferation in epithelial cells via interactions with APC (Ishidate, Matsumine, Toyoshima, & Akiyama, 2000; Matsumine et al., 1996). It is known to be repressed by the transcription factor SNAIL, a driver of EMT (Cavatorta, Giri, Banks, & Gardiol, 2008). Downregulation of *PKP4*, another B-catenin binding protein, (**Figure 2, panel F**) in epithelial-like AsPC1 and PATC153 cells resulted in increased cell proliferation (**Figure 2, panels G, H**) and a significant decrease in cell migration (**Figure 2, panels I, J**). However, in Panc1 cells *PKP4* depletion increased cell migration (**Figure 2, panel K**). *PKP4* encodes Plakophilin 4, a member of the p120-armadillo protein family associated with desmosomes and adherens junctions (Bass-Zubek, Godsel, Delmar, & Green, 2009; Setzer et al., 2004), which also includes p120, encoded by *CTNND1*, a progression driver we identified to be downregulated in advanced PDAC with a significant survival disparity (Mann et al., 2012) that was later validated to be a tumor suppressor in a PDAC mouse model (Reichert et al., 2018). *PKP4* also exhibited a significant survival disparity in PDAC patients (CPH *P*=0.007, **Supplemental Table 3**), suggesting the armadillo proteins play an important role in pancreatic cancer progression and patient survival. PTPRK, a receptor tyrosine phosphatase involved in cell growth and cell adhesion, binds and negatively regulates Beta-catenin (Novellino et al., 2008). Depletion of *PTPRK* (**Figure 3, panel A**) increased cell proliferation of epithelial-like PATC43 and mesenchymal 4039 cells *in vitro* (**Figure 3, panels B and C**) and while PATC43 cells exhibited increased cell migration (**Figure 3, panel D**), 4039 cells exhibited a dramatic decrease in migration (**Figure 3, panel E**) upon *PTPRK* knockdown. This was somewhat surprising, as *PTPRK* was predicted to be a tumor suppressor in advanced PDAC, and 4039 cells are highly mesenchymal-like. To test the effect of *PTPRK* loss *in vivo*, we performed orthotopic injections of 4039 cells into NSG recipient mice containing either a nontargeting shRNA (NTC) or a pool of shRNAs against *PTPRK.* While there was no difference in tumor volume between the two cohorts (**Figure 3, panel F**), we did observe that pancreatic tumors with *PTPRK* depletion were mobile *in vivo*, forming mesenteric masses (**Supplemental Figure 8, panels A and B**) and extensive liver metastases in 3 of 5 mice (**Supplemental Figure 8, panel C**), while control mice did not exhibit liver masses. These data support evidence for *PTPRK* as a tumor suppressor in PDAC which can promote metastases to distal sites. Finally, we investigated the ability of PARD3, an atypical protein kinase belonging to the partitioning defective protein family critical for cell polarization, migration and invasion. PARD3 is involved in cell growth through a protein complex with aPKC and PARD6 at tight junctions and is a negative regulator of Hippo signaling, to promote pancreatic cancer progression. *PARD3* is moderately expressed in mesenchymal cells, and we achieved knockdown in 4039 and Panc1 cells (**Figure 3, panel G**). Panc1 but not 4039 cells exhibited increased growth compared to control cells (**Figure 3, panels H and I**), and while 4039 cells with *PARD3* depletion demonstrated significantly increased migration (**Figure 3, panel J**), the opposite was true in Panc1 cells (**Figure 3, panel K**). *In vivo*, 4039 tumors exhibited significantly increased tumor volumes compared to tumors with the nontargeting shRNA (**Figure 3, panel L**). Collectively, these data support differential roles for *DLG1, PKP4, PTPRK* and *PARD3* in promoting cell proliferation and cell migration *in vitro* in epithelial-like vs. mesenchymal-like PDAC cell lines. *In vivo, PARD3* depletion increased primary tumor growth, while *PTPRK* downregulation promoted metastatic spread, suggesting that downregulation of these two drivers promotes pancreatic cancer progression at different disease stages.

**Figure 2.**
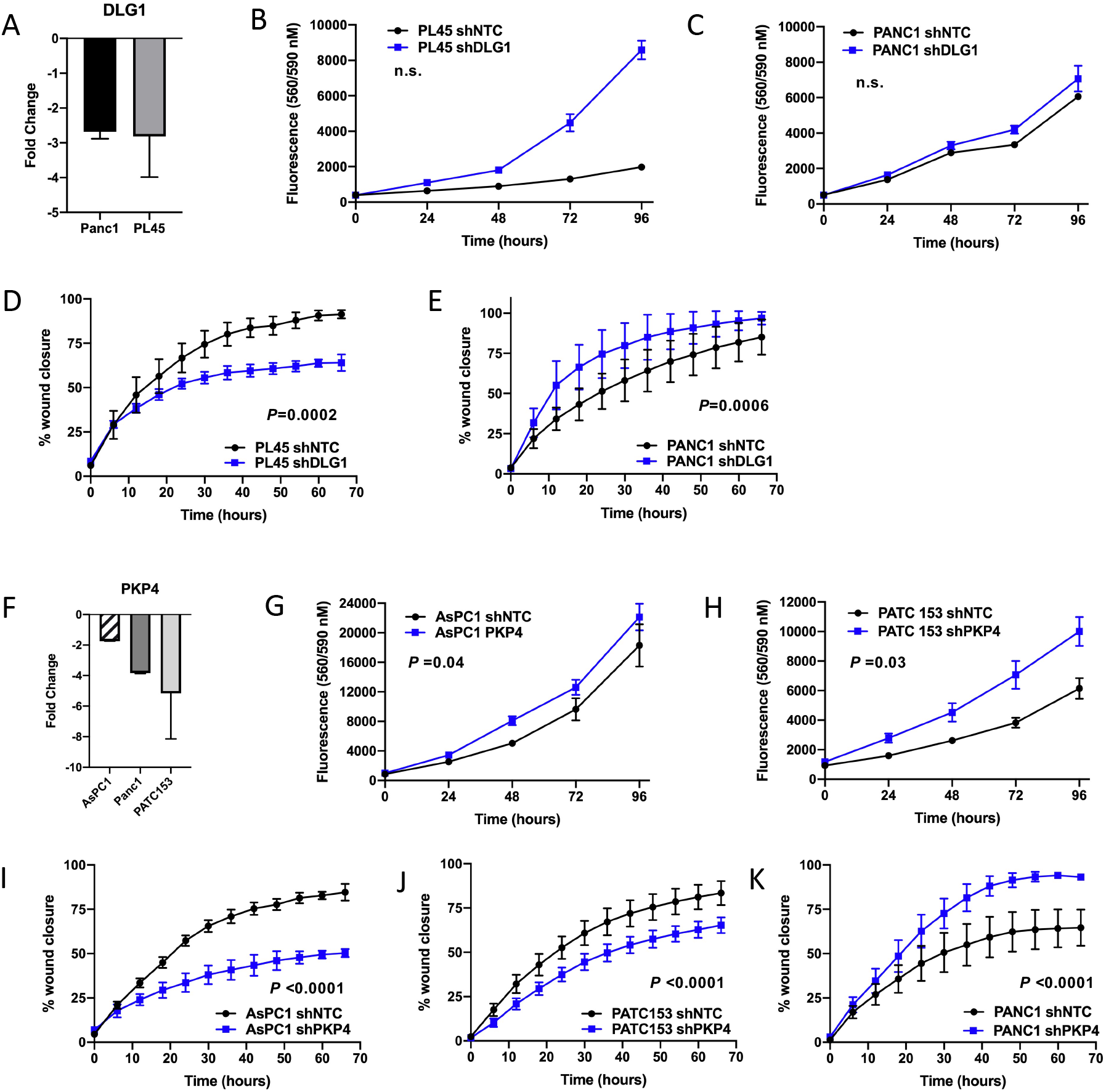
*In vitro* analysis of EPDs in human PDAC cell lines. *DLG1* knockdown using pooled shRNAs (A) induces a trend towards increased cell proliferation in PL45 cells (B) and no change in Panc1 cell proliferation (C). PL45 cells exhibit significantly decreased migration (D, *P*=2E-04) while Panc1 cells show significantly increased cell migration (E, *P*=6E-04). *PKP4* knockdown (F) resulted in increased cell proliferation in both AsPC1 (G, P=0.04) and PATC153 (H, *P*=0.03) cell lines, while both cell lines exhibited significant decreased migratory capabilities (I and J, *P*<0.0001). *PKP4* knockdown in Panc1 cells resulted in significantly increased cell migration (K, *P*<0.0001). Samples were run in triplicate. Statistical analysis was performed using a paired t-test.

**Figure 3.**
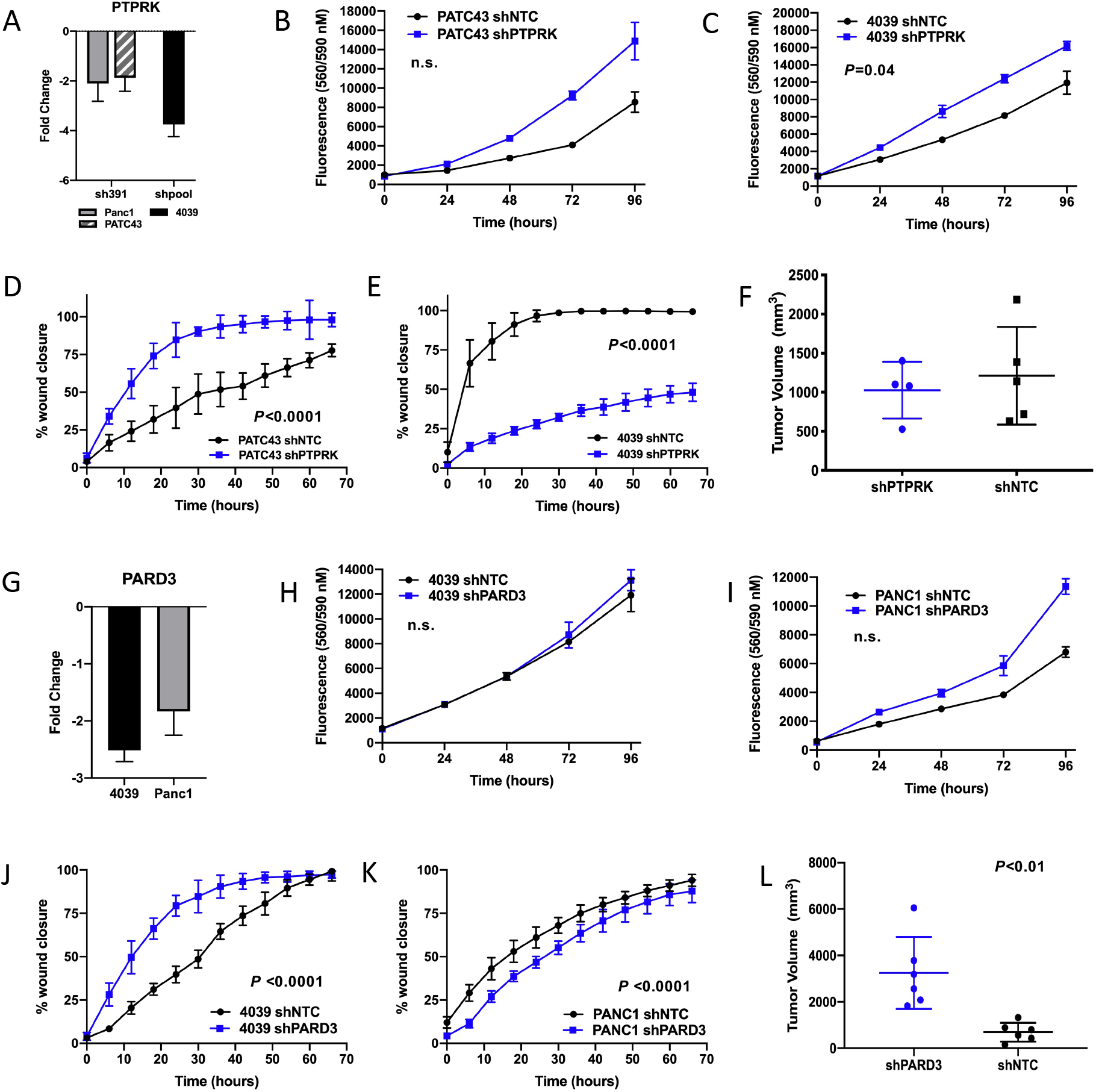
Orthotopic validation of EPDs. Cells stably depleted for EPDs of interest were orthotopically introduced into the pancreas of NSG mice to assess the ability of the predicted tumor suppressor genes to drive tumor formation and disease progression. Individual or pooled shRNAs (A) achieved 2-fold or greater depletion of *PTPRK.* Depletion of *PTPRK* in PATC43 cells using a single shRNA (sh391) showed a trend towards increased proliferation (B) while 4039 cells exhibited a significantly increased proliferative capacity with PTPRK knockdown (C, *P*=0.04). Epithelial PATC43 cells showed significantly increased cellular migration (D, *P*<0.001), while cell migration significantly decreased in 4039 cells (E, *P*<0.001). Orthotopic tumors from 4039 cells with PTPRK depletion showed no change in primary tumor volumes compared to cells with non-targeting shRNA (F). Depletion of *PARD3* (G) using pooled shRNAs in mesenchymal 4039 and Panc1 cells showed no significant change in cell proliferation (H and I). 4039 cells exhibited a significant increase in cell migration (J, *P*<0.0001), while Panc1 cells exhibited a modest but significant decrease in cell migration (K, *P*<0.0001). *PARD3* depletion significantly increased tumor volume in 4039 cells (L, *P*<0.01). Samples were run in triplicate. Statistical analysis was performed using a paired t-test.

### Analysis of recurrent drivers in PDAC metastases

In an effort to elucidate additional tumor cell-intrinsic biology promoting metastases, we next sought to expand our SB Driver Analysis to define selected driver genes in metastatic lesions from the SB_PDAC model. To accomplish this, we first established a new cohort of primary and metastatic lesions from our T2/Onc3 SB_PDAC model and sequenced the SB insertions in matched primary and metastatic lesions using Illumina sequencing. We also re-sequenced our previously reported SB_PDAC tumors and related metastases on the Illumina platform (Mann et al., 2012). SB insertion sequencing data from Illumina and Roche 454 shows high concordance using PCA (Mann et al., 2016). Therefore, we combined these Illumina datasets with the 454 sequencing data from matched primary tumors and related metastases (Mann et al., 2012; Perez-Mancera et al., 2012) to establish a population of 31 metastatic primary tumors, driven by four independent SB transposons, and 79 matched metastatic lesions, including 46 liver lesions. SB Driver Analysis was performed using mapped insertions with 3 reads or higher from 454 sequencing data and 10 or greater reads from Illumina sequencing data, based on our previously established criteria for analyzing low complexity SB sequencing data.

329 progression drivers recurrent in three or more tumors were defined from the 31 metastatic primary tumors (**Supplemental Table 5**). Sixty percent of the drivers overlapped with the progression drivers we defined from the larger cohort analysis of 172 primary tumors (Newberg, Mann, et al., 2018), showing conservation of these recurrent drivers in metastatic and non-metastatic primary tumors. Further, 50 genes were defined as EPDs (**Supplemental Table 1**). Driver analysis using SB insertions from 79 metastatic lesions isolated from the liver, lung, diaphragm, peritoneum, stomach and lymph nodes defined 59 genome-wide significant (familywise error rate, FWER) trunk drivers highlighted in the oncoprint shown in **Figure 4, panel A and Supplemental Table 5**. We manually removed *Xpo4* and *Armc3* from the analysis output as they contained insertions within the promoter region that also mapped within a neighboring gene. The majority of drivers were defined from metastatic lesions across different organ sites from multiple animals, suggesting they are common to the metastatic process (**Supplemental Table 5**). For some animals, the shared clonal origins, or identity by decent, was evident for multiple lesions based upon shared nucleotide insertions (**Supplemental Figure 9, panel A**). Further, shared insertions between primary and metastatic lesions were evident for a subset of the cohort (**Supplemental Figure 9, panel B**). Interestingly, only 19 of the 59 metastatic drivers overlapped with the defined progression drivers from the 31 matched primary tumors, suggesting that the majority of SB insertions in the primary drivers were not selectively maintained in the metastatic lesions and that new drivers were acquired during the metastatic process. *Foxp1* and *Setd5* were the only two EPDs in common between related primary (progression drivers) and metastatic lesions (trunk drivers) (**Supplemental Table 1**). *SETD5* upregulation has recently been linked to MEK resistance in PDAC (Z. Wang et al., 2020). The most recurrent driver, *Foxp1*, was present in 26 of the metastatic lesions from liver, lung and lymph nodes, suggesting that this driver plays a role in both establishing and promoting pancreatic cancer progression in the SB model. Interesting, only four of the 31 matched primaries had detected insertions in *Foxp1.* This low rate of concordance may be due to a failure to capture sub-clonal populations in the primary tumors containing *Foxp1* insertions. Alternatively, selection of *Foxp1* insertions may have occurred during the metastatic process. Next, we interrogated the expression of the metastatic drivers using available RNASeq data from liver metastases (EGAD0000003584). This dataset included 11 primary pancreas tumors and 34 unrelated liver metastases. We derived a metagene for the expression of the human homologs of the 83 FDR-significant metastasis drivers for each cohort and found that the mean expression was significantly lower in the liver metastases compared to the primary tumors (**Figure 4 panel B**). These data support the role for these drivers as tumor suppressors in human PDAC liver metastases.

**Figure 4.**
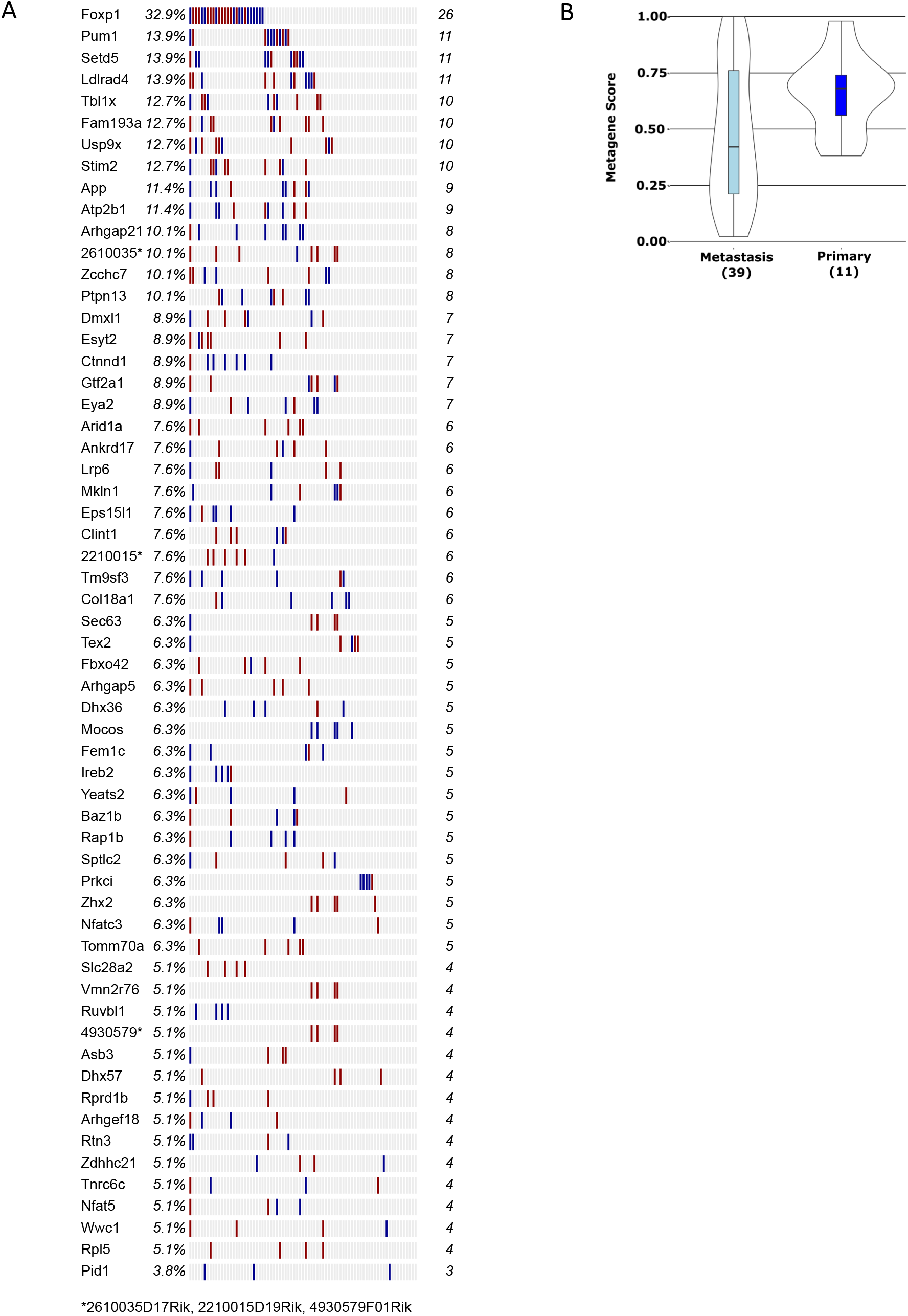
Driver enrichment analysis in metastatic lesions. Driver analysis was performed for 74 metastatic lesions isolated from liver, lung, diaphragm and lymph nodes for which matching primary tumor sequence information was available. The oncoprint (A) shows the recurrent drivers statistically significant after FWER multiple-testing correction. The gene name and percentage of metastatic lesions containing insertions in each driver on the left, with the number of tumors on the right. Blue lines indicate insertions in the sense direction, while red lines indicate insertions in the antisense direction with respect to the gene coding region. (B) An expression metagene derived using the human homologs of the SB_PDAC metastatic drivers shows significantly decreased expression in human PDAC liver metastases compared to primary tumors (*P*=0.05).

Because the majority of the metastases we collected from the SB_PDAC model were derived from liver, we further explored whether we could uncover liver-specific drivers that may lend insight to the biology underlying pancreas liver metastases. We statistically defined 55 FWER (genome-wide)-significant trunk drivers from 46 liver metastases isolated from 15 mice (**Supplemental Table 6**). The majority (48) from this “liver only” driver analysis overlapped with drivers defined from the all-metastasis (“all mets”) analysis but were not defined as drivers in the matched primary analysis. Seven drivers were identified as EPDs: *Foxp1, Setd5, Dock9, Lpp, Mkln1, Pkp4*, which we functionally validate in this manuscript, and *Rbms1.* The insertion patterns for these 7 drivers in metastatic lesions are shown in **Supplemental Figure 10**. The overall insertion patterns in these genes suggest that these drivers are downregulated by SB. Insertions across the *Foxp1* coding sequence showed distinct patterns. Some related metastases showed maintenance of an SB insertion at the same nucleotide address, suggesting identity by descent from a founding clone with this *Foxp1* insertion, while other related metastases showed insertions across the coding region, suggesting either acquired insertions in individual metastatic clones through convergent evolution or by mobilization of an insertion from a founding clone and reintegration at unique sites throughout the *Foxp1* coding region. Clustered insertions in the sense direction may drive a truncated *Foxp1* transcript, as previously hypothesized by Rad *et al.* (Rad et al., 2015). Taqman analysis showed *Foxp1* expression was highly variable in primary tumors and did not correlate with the presence or directionality of SB insertions (**Supplemental Figure 10, panel H**). Five related metastatic liver lesions with SB insertions in *Foxp1* had a lower mean expression level than the primary tumor samples. *Lpp* encodes a protein involved in cell adhesion that was recently implicated in a model of breast cancer lung metastasis (Ngan et al., 2017). *Tbx1l*, a liver-specific metastatic trunk driver, encodes a WD40-containing protein implicated in regulation of Wnt and NFKB signaling in hepatocellular carcinoma and colon cancer (Qu et al., 2016; Zhang et al., 2016). This gene was not identified in primary PDAC tumors, and the insertion pattern in liver metastases suggests that the coding region is deregulated. Seven liver-specific drivers, including *Mocos and Tomm70a*, were private to individuals, while eight (*Capn7, Naa15, Camk2d, Thsd4, Fut8, Utrn, Ophn1, Mid1*) were defined from liver lesions from multiple individuals. *Camk2d* encodes a kinase involved in calcium signaling, while *Ophn1* encodes a protein involved in GTP hydrolysis of Rho proteins, including RHOA. Collectively, these drivers of liver metastases are involved in a variety of cellular processes, including amino acid metabolism, calcium signaling, Wnt signaling and transcriptional regulation. Further, thirteen of the drivers we identified in this analysis were also drivers in an SB model of HCC (Newberg, Mann, et al., 2018), suggesting that the selection of insertions in these genes may be driven by the microenvironment.

### Clonal origins of primary PDAC tumors and related metastases

From our bulk tumor sequence analysis of pancreas tumors, we observed a large degree of inter- and intra-tumor heterogeneity based on the number of insertions per tumor, the sequencing depth of insertions and the profile of recurrent genes in each tumor. To further characterize the intratumor heterogeneity and explore clonal relationships, we performed regional sequencing on 10 individual tumors from our expanded T2/Onc3 tumor cohort. Pancreas masses were sub-sampled at necropsy into 3 to 5 equally sized regions, from pancreas head to tail, while collecting an adjacent tumor region for histological confirmation of PDAC and any observed metastases (**Figure 5, panel A**). Each region was sequenced for SB insertions and analyzed for clonal relationships with neighboring regions using SB transposon insertions as markers of evolutionary history. This analysis was possible using the combination of the low copy T2/Onc3 transposon (~30-40 present in each cell) and the use of the inducible Rosa26-LSL-SBase (Dupuy et al., 2009), whose activity controls the number of transposition occurrences at a frequency that maintains both clonal and sub-clonal insertions. We found identical insertions shared across regions, independent insertions in genes recurrently hit across regions and genes uniquely selected in sub regions of a single tumor. For tumor Malt 285, the presence of insertions in *Ankrd52, Cluh* and *Golph3* occurring at the same nucleotide in all three tumor sub regions (**Figure 5, panel B)** suggests identity by descent for a sub clonal population, whereby these early selected events in a precursor clone persisted through clonal expansion, either through positive or neutral selection. The increased read depth of these insertions in Regions A and D compared to Region C suggests clonal expansion of cells containing these events in these two regions. Single-cell sequencing analysis of SB insertions in 28 cells isolated from a low passage cell line established from the M285 pancreas tumor revealed co-occurrence of the clonal insertions in *Cluh* and *Golph3* in two cells (M285-76 and M285-15, **Figure 5, panels C and D**), in addition to other insertions common to subpopulations of cells shown by hierarchical clustering. From the bulk tumor analysis, Region A contained 12 EPDs, while Region C contained 16 and Region D contained 6. *Taf1* was common in all regions, while insertions in *Foxp1* and *Arhgef12* were found in Regions A and C. Similar observations of shared insertions and shared genes were made from regional sequencing of nine additional tumors. For Malt 299 primary tumor regions, 11 genes were in common across all three regions and 93 additional genes shared by two regions (**Supplemental Figure 11, panel A**). While these genes are in common, they do not share common insertions across tumor regions. This suggests that there is positive selection for these genes, and by regionally sequencing the end-stage tumor, we are sampling multiple insertion sites in selected genes that represent either remobilization and re-integration of founder insertions across the coding region of these genes or independent integration and selection of insertions through convergent evolution in a sub-clonal manner. These two possibilities cannot be distinguished here.

**Figure 5.**
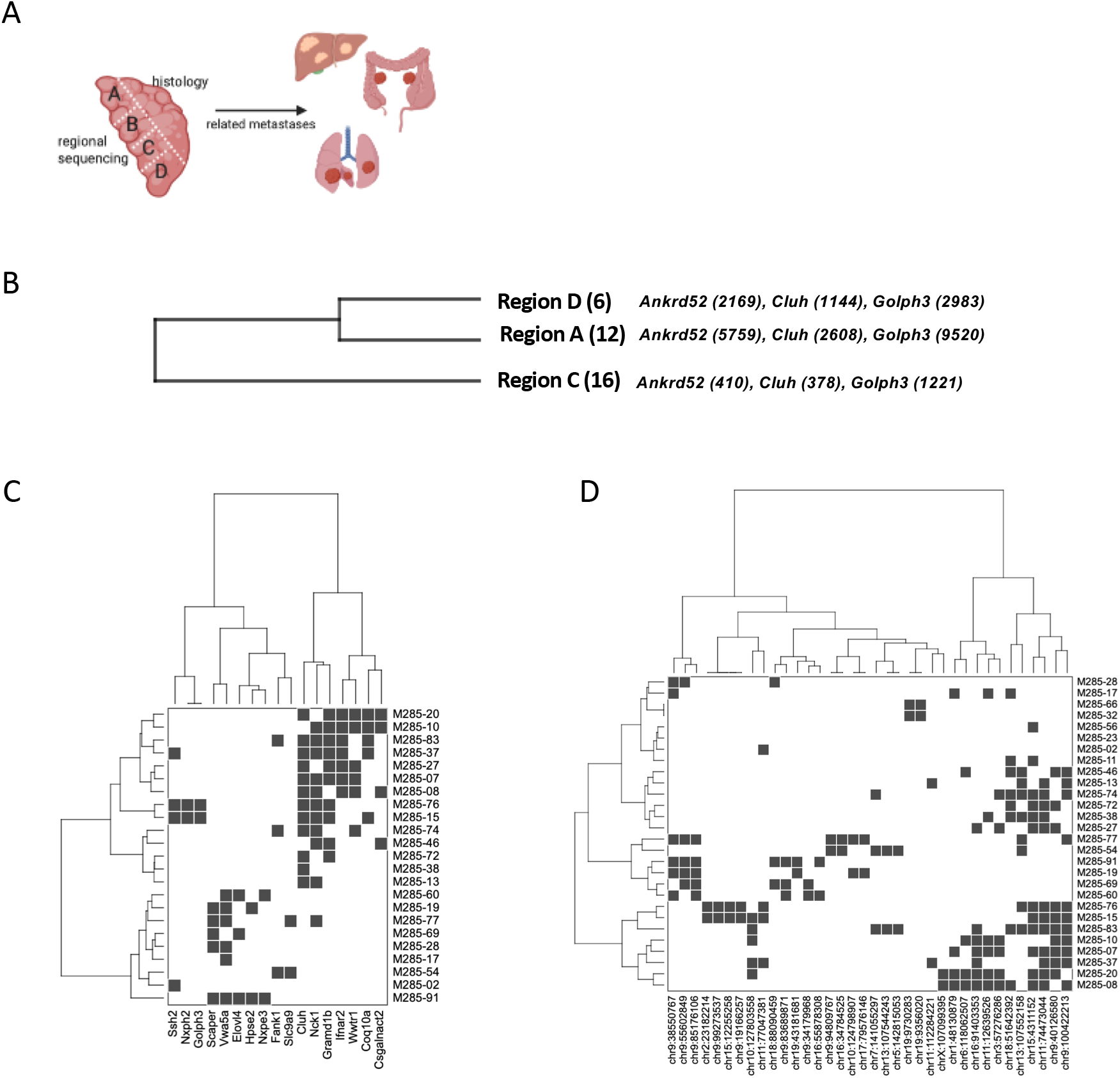
Tracing clonal relationships in primary tumors using regional and single-cell sequencing of SB insertions. Pancreas tumors from the T2/Onc3 SB_PDAC model were regionally dissected (A) and histologically confirmed. Related metastases isolated from the liver, mesentery and lung were snap frozen for future analyses. Regional sequencing of a primary tumor uncovered clonal relationships between regions shown by hierarchical clustering of coincident genes with SB insertions (B). The number of EPDs identified in each region is shown in parentheses. All three regions share 3 insertions at identical nucleotides in *Ankrd52, Cluh* and *Golph3.* The number of reads per insertion in each gene is shown. Single cell analysis of SB insertions present in 28 cells isolated from a cell line established from Malt 285 primary tumor shows coincident genes (C) and SB insertion addresses (D) that indicate identity by descent for subpopulations of cells. Insertions in *Cluh* and *Golph3* found by bulk tumor sequencing co-occur in cells M285-15 and M285-76.

We next expanded our analyses to define clonal relationships among metastatic lesions and between metastases and the related primary tumor using SB insertion profiles. Comparing regional sequencing insertion profiles from primary tumor Malt 299 with its 5 related liver metastases failed to detect shared insertion sites present at the same nucleotide. This could be attributed to an insufficient depth of sequencing in the primary regions to trace the clonal origin of the metastatic lesions, or local hopping and selection of reintegrated transposons at different TA dinucleotides across the coding region in primary tumors and/or related metastases. Regardless, we could define clonal relationships among liver metastases. SB insertions in *Foxp1* and *Fam83b* were identified at the same nucleotide in all five related liver metastases, suggesting these liver metastases are derived from the same tumor clone (**Figure 6, panel A and Supplemental Figure 17, panel B**). The number of total SB insertions in each liver metastasis (MS) inclusive of the genes with shared insertion addresses is shown in parentheses (**Figure 6, panel A**). MS2 and MS5 are distinguished both by the number of insertions and the acquired insertion in *Dopey1* in MS2. The insertions in the clonal markers *Foxp1, Fam83b* and *Dopey1* are retained by MS2 at high sequencing read depths, suggestive of clonal expansion. The sub clonal origin of MS8 and MS3 is marked by the acquired shared insertion in *Dync1li1*, while most of the insertions in these two lesions are distinct. Together, these data demonstrate the clonal origins of these liver metastases that continue to evolve independently. Importantly, single cell sequencing analysis of SB insertions in 22 cells isolated from a low passage cell line established from the M299 pancreas tumor showed co-occurrence of the clonal insertions in *Foxp1* and *Fam38b* at the same nucleotide positions mapped in the liver lesions in 18 and 14 cells, respectively (**Figure 6, panels B and C**). Ten cells showed coincident insertions in both genes, confirming that these insertion events were selected and maintained in the primary tumor sub clone that gave rise to the liver metastases. For tumor Malt 53, a clonal insertion in *Foxp1* present in pancreas Regions 1 and 5 from the primary tumor (**Supplemental Figure 11, panel C**) was maintained in all 3 lung metastases from different lobes (**Figure 6, panel D and Supplemental Figure 11, panel D**). The sub-clonal origin of LU_IL and LU_ML is demonstrated by 3 shared insertions in *Tbcc, Cdc20b* and at chr1:91223279, yet these two lesions are distinct from each other based on the number of unique SB insertions (number in parentheses) and the distinct genes hit. LU_ML acquired the greatest number of insertions, suggesting clonal diversion and expansion from LU_IL. Taken together, these results highlight the underlying intratumor heterogeneity of primary tumors and related metastases and demonstrate that metastatic lesions often share clonal origins with a distinct but minor sub clone of the primary tumor and acquire additional highly selective insertions in their distal microenvironment.

**Figure 6.**
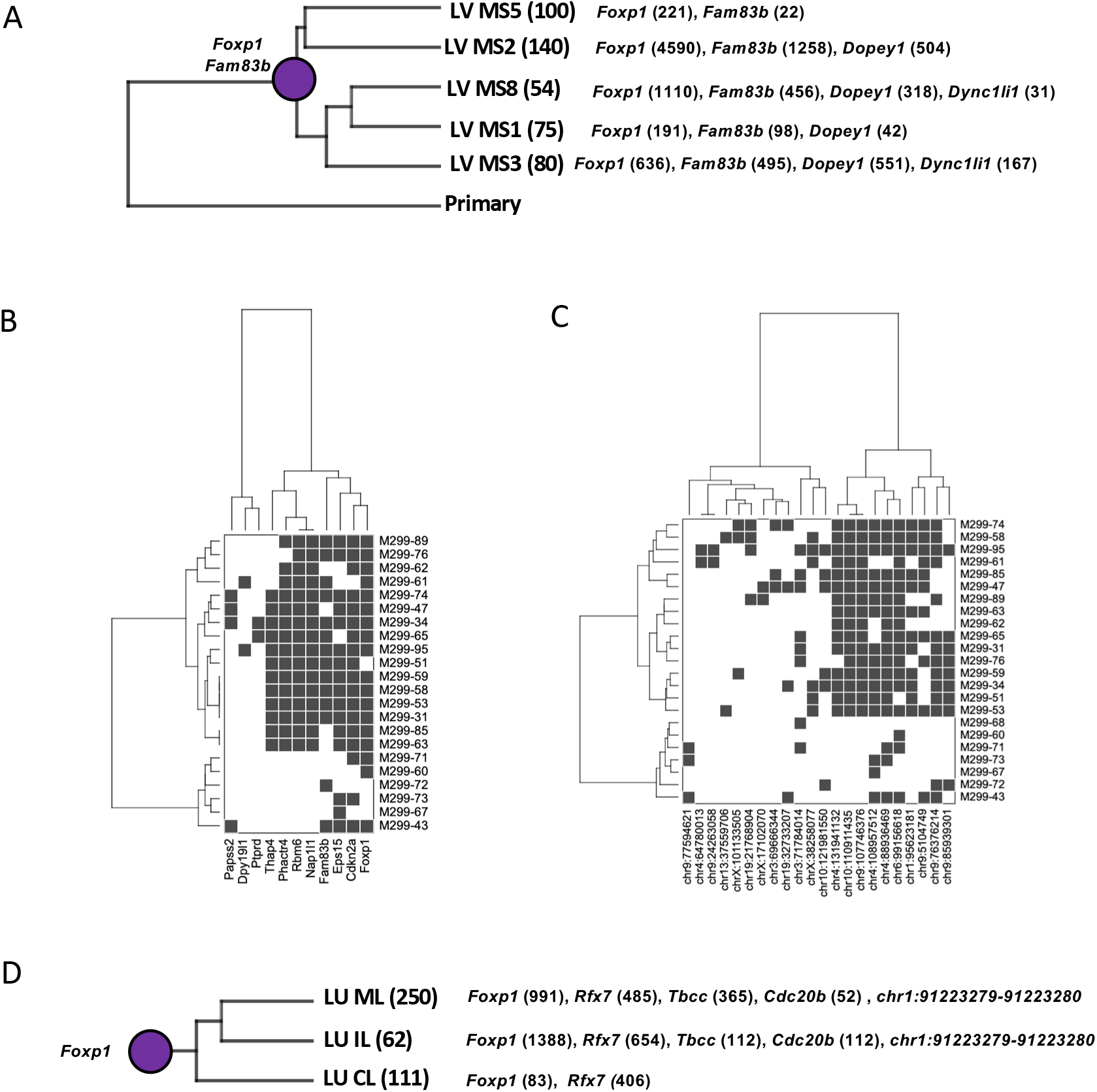
Evolutionary analysis of primary tumors and related metastases. Sequence analysis of SB insertions from 5 related liver nodules (A) defined clonal relationships by hierarchical clustering based on coincident genes with SB insertions. The number of unique insertions in each liver nodule is indicated in parentheses adjacent to the mass designation. All five lesions show identity by descent based on the presence of SB insertions maintained at fixed nucleotide positions in *Foxp1* and *Fam83b* originating from an unknown (primary) sub-clone (purple circle). The read depth for shared insertions is indicated in parentheses next to the gene name. MS5 is distinct from all other masses as it lacks a fixed insertion in *Dopey1.* MS3 and MS8 share an acquired 4^th^ fixed SB insertion in *Dync1li1*. Single cell analysis of SB insertions present in 22 cells isolated from a cell line established from Malt 299 primary tumor shows coincident genes (B) for the majority of the tumor cells and uncovered the primary tumor origin of the shared insertions in *Foxp1* and *Fam83b* present in the liver metastases (C). Sequencing of three metastatic masses from different lung lobes in an individual animal identified fixed insertions in *Foxp1* from the primary tumor and an acquired insertion in *Rfx7.* LU_IL and LU_ML are distinguished from LU_CL by three additional shared insertions in two genes and in one intergenic region. LU_IL and LU_ML are distinguished by the number of unique insertions (in parentheses) and by the genes with SB insertions.

## DISCUSSION

Identifying drivers of metastasis is an important unmet need in pancreatic cancer research and holds potential to impact patient diagnosis, predict disease recurrence and potentially uncover new therapeutic opportunities. Yet, the molecular heterogeneity present in pancreatic cancers has confounded efforts to identify novel, recurrent cooperating drivers in metastatic disease. Rather, studies to date have reinforced the importance of long-known recurrent drivers KRAS, TP53 and SMAD4. Two Sleeping Beauty screens in mouse models of pancreatic cancer (Mann et al., 2016; Mann et al., 2012; Perez-Mancera et al., 2012) identified over one-thousand genetic drivers of disease that cooperate with oncogenic *Kras*^G12D^, most with little known about their potential role in pancreatic disease initiation and development. In this study, we have leveraged our Sleeping Beauty mouse model of pancreatic cancer and our statistical SB Driver Analysis method to define early recurrent cooperating drivers in primary tumors that promote disease progression and, for the first time, define selected drivers in metastatic lesions. Our EPD analysis reaffirmed the importance of 79 highly recurrent and clonal drivers previously reported in the top 150 most frequent progression drivers (Newberg, Mann, et al., 2018), including *Dlg1, Pard3, Pkp4* and *Ptprk* that we functionally validated in this manuscript, and prioritized 36 new drivers, including *Ep300* and *Smad3*, which were identified as two nodes in the EPD protein interaction network. Indeed, we showed that low expression of an EPD metagene was associated with statistically significantly reduced disease-free survival in human pancreatic cancer patients. Further, these same genes showed a significantly reduced expression in liver metastases compared to primary tumors in human patients, suggesting that the mis-regulation of these drivers plays a role not only in primary tumor progression but also potentially in the initiation and maintenance of liver metastasis, the most frequent metastatic site in pancreatic cancer.

To address the clonal origins of drivers in pancreatic tumors, we performed the first in-depth analysis of the inter- and intra-tumoral heterogeneity present in the SB_PDAC mouse model using regional bulk and single-cell sequencing of SB insertions in individual primary tumors, defining clonally selected genetic events common to all regions as well as sub-clonal and private insertions. Further, by sequencing primary tumors and related metastases, we showed using SB insertion profiles that primary and metastatic tumors from an individual are more related to each other than to tumors from unrelated individuals. This observation has been reported in human pancreatic tumors (Campbell et al., 2010; Yachida et al., 2010) and more recently in a KPC mouse model (Niknafs et al., 2019). Here, using insertion nucleotide sharing as a marker of evolutionary history and read-depth as a proxy for clonal expansion, we uncovered the primary tumor sub clonal origins of metastases and defined the insertional heterogeneity that distinguishes related metastatic lesions.

Defining drivers promoting metastasis to characterize underlying molecular vulnerabilities is of paramount importance for improving patient outcomes in pancreatic cancer. To this end, we defined 59 drivers enriched in metastases from the SB_PDAC model and human homologs of these metastasis drivers showed significantly lower expression in liver metastases compared to primary tumors, strongly supporting the important roles of these genes as tumor suppressors in the metastatic process. Further, we identified 16 liver-specific metastasis drivers not present in PDAC metastases from other organ sites. Indeed, several of these liver metastasis drivers also have biological relevance to an SB model of hepatocellular carcinoma (Newberg, Mann, et al., 2018), providing evidence that the microenvironment partly influences the selection of insertions while highlighting biological commonalities between HCC and liver metastases. *PKP4* was defined as both an EPD and a liver-specific trunk driver and we functionally validated this driver in both human patient samples and in human PDAC cells. *PKP4* was previously functionally validated as a tumor suppressor in triple negative breast cancer cells that accelerates tumor growth (Rangel et al., 2016). Thus, the recurrence of drivers in the SB_PDAC model can lend insight into the underlying genetic and epigenetic mechanisms of pancreatic cancer metastases that have proven difficult to elucidate from human sequencing analysis. Our preliminary analyses of tumor heterogeneity in primary tumors and related metastases underscore the underlying drivers common to individual disease, the clonal origins of metastases and their unique molecular signatures which may have important implications for defining novel molecular signatures common to both primary and metastatic lesions and for tracing the development of early metastasis, which is a significant focus in the field.

Genetic screens are powerful approaches to identify both known and novel relationships between genes and genetic pathways that cooperate to promote disease. This is the first study to report and functionally validate novel metastatic drivers identified in a *Sleeping Beauty* genetic screen with direct patient relevance. One important advantage of the SB system for pancreatic cancer gene discovery is the fact that low tumor cellularity can be overcome by enriching for cells that have mobilized transposons. Leveraging our unique SB_PDAC mouse model, we can investigate the underlying differences in transcriptional, biochemical and metabolic signatures between primary tumors and related metastases to experimentally define druggable vulnerabilities of metastatic disease. Indeed, we have generated stable cell lines from metastatic SB_PDAC primary tumors which retain their ability to metastasize in B6 recipient mice. This unique transplantable model with SB insertion-based intratumor heterogeneity as a salient feature will enable future studies that trace the clonal evolution of pancreas disease during development and in response to drug administration in an immune competent setting and can further be harnessed to define innate and acquired routes of resistance. Our current study has laid the foundation for defining metastasis drivers and provides a complementary guide to ongoing studies in human pancreatic cancer patients to combat metastatic disease.

## MATERIALS and METHODS

### Cell Lines and in vitro assays

Human pancreatic cancer cell lines were purchased from ATCC and grown under recommended media conditions. Human PATC cell lines were grown in RPMI as previously described (Kang et al., 2015). Cell proliferation was determined using Cell Titer-Blue Viability assays in triplicate (Promega G8080) in triplicate. Wound healing assays were performed using IncuCyte Zoom following the supplier’s experimental guidelines. Cells were synchronized using mitomycin-C (Sigma Aldrich) at 5-10ug/ml concentration for 2 hours prior to seeding in triplicate on 96-well ImageLock plates (Essen BioScience) and grown to 95% confluency. Scratch wounds were created with a WoundMaker (Essen BioScience) and media was replaced after a PBS wash. Images of cell migration/wound healing were recorded every 6 hours for a duration of 96-120 hours using the IncuCyte Kinetic Cell Imaging System and analyzed with IncuCyte Zoom software. Statistical analysis was performed using GraphPad prism software using a paired t-test to compare knockdown cells to matched controls with a non-targeting shRNA.

### RNA interference and expression analysis

shRNAs were obtained as viral particles from Dharmacon (GE Biosciences). Cells were transduced and stably selected under puromycin prior to molecular analysis. For qPCR analysis, total RNA was extracted using EZNA total RNA kit (R6834-02, Omega Bio-Tek). cDNA synthesis was performed using qScript® cDNA Synthesis Kit (95047-100, Quanta Bio) and qPCR analysis was performed using Sybr Green with primer sequences obtained from PrimerBank (X. Wang, Spandidos, Wang, & Seed, 2012). Taqman probes for *Foxp1* and *Gapdh* were purchased from Life Technologies. For expression analysis using Nanostring or RNASeq, total RNA was extracted using Qiagen RNeasy mini kits. Nanostring was run on two different passages for each cell line and was analyzed using IRON (Welsh, Eschrich, Berglund, & Fenstermacher, 2013).

### Antibodies and western blotting

Cells were lysed in RIPA lysis buffer (100 mM Tris pH7.5, 300mM NaCl, 2mM EDTA, 2% NP-40, 2% Sodium deoxycholate and 0.2% SDS) with 1% protease inhibitor (Amresco M250-1ML). Cell lysates were sonicated using Diagenode Bioruptor Plus for 15 total minutes (30 seconds on and 30 seconds off). Total protein concentration was measured using a BCA assay kit (23227 Pierce). Immunoblotting was performed using 50μg of protein run on 8% SDS-PAGE gels and transferred to nitrocellulose membranes. Membranes were blocked for one hour using Odyssey Blocking Buffer (Cat.No.927-50000, Li-COR Biosciences). Antibodies: Vimentin (5741S), E-Cadherin (3195S), GAPDH (rabbit, 2118S) from Cell Signaling and anti-rabbit IgG, 926-32211 secondary from LI-COR Biosciences. Blots were imaged using Li-COR Odyssey Fc Imaging System.

### Mice

Human Panc1 (1×10^5) and 4039 (5×10^4) cells were orthotopically introduced into the pancreas of male and female NSG recipient mice (JAX 005557) according to methods described previously (Kim et al., 2009). Isogenic cell lines (knockdown vs. control) were matched for cell passage, implanted into littermates and aged concurrently. Mice were aged for 45 days and then euthanized to measure primary tumor growth and metastatic incidence. All procedures were conducted under approved IACUC protocols.

### Sequencing of SB insertions and SBCapSeq driver analysis

Sequencing SB insertions using Roche 454 sequencing was described previously (Mann et al., 2012). Data processing and driver analysis were performed using SBCapSeq Driver Analysis pipeline (Newberg, Black, et al., 2018) with the following modifications. To define early progression drivers (EPDs) using 454 sequencing data, bed files were filtered to remove all insertions with fewer than 3 sequencing reads prior to statistical analysis. Only drivers identified in 3 or more independent tumors (5% of cohort) were considered recurrent EPDs. For analysis of metastatic drivers, insertion data from 454 and Illumina HiSeq were combined (as described in (Mann et al., 2016)) and the analysis was run using sequencing reads of 3 or higher from 454 data and 10 or greater reads from Illumina. Single cell sequencing was performed as described previously (Mann et al., 2016). 100 DAPI-positive single cells were FACS-sorted from early-passage cell lines (P2) established from primary tumor explant cultures. Onc excision PCR was used to confirm the presence of mobilized transposon from the donor site, confirming the tumor-origin of the cells. Two-dimensional hierarchical clustering of conserved SB insertions or recurrent genes containing SB insertions was performed in Python (2.7.10) with the scipy toolbox using the Hamming distance metric with Ward’s linkage method of agglomeration.

### Mutation analysis using human PDAC datasets

Mutation data from three pancreatic cancer cohorts was downloaded from cBioPortal (Cerami et al., 2012). Data sets are denoted as per cBioportal coding, based on facility or project contributing the data: QCMG (Queensland Centre for Medical Genomics, n=383) (Bailey et al., 2016), TCGA (The Cancer Genome Atlas, n=150) (Network, 2017), UTSW (University of Texas Southwestern, n=109) (Witkiewicz et al., 2015). GOseq analysis (Young, Wakefield, Smyth, & Oshlack, 2010) was performed using both Gene Ontology terms (The Gene Ontology, 2019) and also using a set of gene lists downloaded from the Enrichr website (Kuleshov et al., 2016). Gene lists were downloaded so that gene length bias could also be accounted for via GOseq in the analyses based on the Enrichr lists. In each analysis, gene sets were considered to be significantly enriched if their FDR adjusted p-value was less than 0.05.

### Gene expression and survival analyses using human PDAC datasets

RNA-seq data and clinical information were downloaded from the International Cancer Genome Consortium (Zhang et al., 2019) data portal for a pancreatic cancer cohort of 90 patients (PACA-AU: Pancreatic Cancer - Australia, release 28), 84 for whom both RNA-seq and survival data were available (Bailey et al., 2016). RNA-seq read counts (per gene) were normalized using the workflow outlined in the limma software package for R (Law, Chen, Shi, & Smyth, 2014), and singular value decomposition was used to generate an expression-based metagene from the genes of interest. The log-rank test was then used to determine whether high versus low values of the metagene (top 50% of values versus bottom 50%) were associated with differences in patient survival characteristics.

### EGA dataset for primary vs. liver metastasis

Access was obtained for the dataset “Genomic Profiling of Advanced Pancreatic Cancer to inform therapy - RNA-Seq mapped reads” (EGAD00001003584) via the European Genome-Phenome Archive. These data consist of BWA-mapped RNA-seq reads (reference: hg19) for 50 pancreatic cancer tumours (11 primary tumours, and 39 metastatic tumours) from the COMPASS trial (Aung et al., 2018). The featureCounts tool from the Subread software package was used to generate mapped fragment counts per gene (Liao, Smyth, & Shi, 2014) based on the aligned BAM data. These per-gene read counts were normalized using the workflow outlined in the limma software package for R (Law et al., 2014), and singular value decomposition was used to generate expression-based metagenes to summarize the activity of specific gene sets of interest.

## Accessions

NCBI BioProject accessions: PRJNA664792 for single-cell-SBCapSeq data; PRJNA664797 for Illumina SplinkSeq.

## Supplemental Figures

Available from Figshare http://dx.doi.org/10.6084/m9.figshare.13017830

## Supplemental Tables

Available from Figshare http://dx.doi.org/10.6084/m9.figshare.13017818

## Author contributions

K.M.M. designed and supervised the study and drafted the manuscript; M.M. and S.G. performed gene knockdown studies; S.G performed proliferation and migration assays; K.M.M., S.G. and D.J.J. performed orthotopic surgeries; J.L.M provided pathology expertise; D.J.J. and H.L. made genomic libraries for DNA sequencing; J.Y.N., M.B.M and M.A.B performed bioinformatic analyses. All authors read and approved the final manuscript.

## Acknowledgements

The authors thank Jason Fleming, Moffitt Cancer Center, for human PATC cell lines and Neal Copeland and Nancy Jenkins, MD Anderson Cancer Center, for early support of this work. We are indebted to the expertise of Sean Yoder and the Moffitt Cancer Center Molecular Genomics Core, Eric Welsh and the Moffitt Cancer Center Biostatistics and Bioinformatics Core, Joe Johnson and the Moffitt Cancer Center Microscopy Core, and Noel Clark and the Moffitt Cancer Center Tissue Core supported by CCSG P30-CA076292. We thank Bethanie Gore, Margi Baldwin, Cris Rivera and the University of South Florida Comparative Medicine facility for assistance with *in vivo* studies and Qing Yan, Tiffany Arnold and Julia Billington for technical assistance. Finally, we thank Marco Napoli and Elsa Flores, Moffitt Cancer Center, for their generous support and expertise with Incucyte assays.

## REFERENCES

Aung, K. L., Fischer, S. E., Denroche, R. E., Jang, G. H., Dodd, A., Creighton, S., … Knox, J. J. (2018). Genomics-Driven Precision Medicine for Advanced Pancreatic Cancer: Early Results from the COMPASS Trial. Clinical cancer research: an official journal of the American Association for Cancer Research, 24(6), 1344–1354. doi:10.1158/1078-0432.CCR-17-2994

Bailey, P., Chang, D. K., Nones, K., Johns, A. L., Patch, A. M., Gingras, M. C., … Grimmond, S. M. (2016). Genomic analyses identify molecular subtypes of pancreatic cancer. Nature, 531(7592), 47–52. doi:10.1038/nature16965

Bass-Zubek, A. E., Godsel, L. M., Delmar, M., & Green, K. J. (2009). Plakophilins: multifunctional scaffolds for adhesion and signaling. Curr Opin Cell Biol, 21(5), 708–716. doi:10.1016/j.ceb.2009.07.002

Biankin, A. V., Waddell, N., Kassahn, K. S., Gingras, M. C., Muthuswamy, L. B., Johns, A. L., … Grimmond, S. M. (2012). Pancreatic cancer genomes reveal aberrations in axon guidance pathway genes. Nature, 491(7424), 399–405. doi:10.1038/nature11547

Bonello, T. T., & Peifer, M. (2019). Scribble: A master scaffold in polarity, adhesion, synaptogenesis, and proliferation. The Journal of cell biology, 218(3), 742–756. doi:10.1083/jcb.201810103

Campbell, P. J., Yachida, S., Mudie, L. J., Stephens, P. J., Pleasance, E. D., Stebbings, L. A., … Futreal, P. A. (2010). The patterns and dynamics of genomic instability in metastatic pancreatic cancer. Nature, 467(7319), 1109–1113. doi:10.1038/nature09460

Cavatorta, A. L., Giri, A. A., Banks, L., & Gardiol, D. (2008). Cloning and functional analysis of the promoter region of the human Disc large gene. Gene, 424(1-2), 87–95. doi:10.1016/j.gene.2008.07.040

Cerami, E., Gao, J., Dogrusoz, U., Gross, B. E., Sumer, S. O., Aksoy, B. A., … Schultz, N. (2012). The cBio cancer genomics portal: an open platform for exploring multidimensional cancer genomics data. Cancer discovery, 2(5), 401–404. doi:10.1158/2159-8290.CD-12-0095

Collisson, E. A., Sadanandam, A., Olson, P., Gibb, W. J., Truitt, M., Gu, S., … Gray, J. W. (2011). Subtypes of pancreatic ductal adenocarcinoma and their differing responses to therapy. Nature medicine, 17(4), 500–503. doi:10.1038/nm.2344

Connor, A. A., Denroche, R. E., Jang, G. H., Lemire, M., Zhang, A., Chan-Seng-Yue, M., … Gallinger, S. (2019). Integration of Genomic and Transcriptional Features in Pancreatic Cancer Reveals Increased Cell Cycle Progression in Metastases. Cancer cell, 35(2), 267–282 e267. doi:10.1016/j.ccell.2018.12.010

Dupuy, A. J., Rogers, L. M., Kim, J., Nannapaneni, K., Starr, T. K., Liu, P., … Copeland, N. G. (2009). A modified sleeping beauty transposon system that can be used to model a wide variety of human cancers in mice. Cancer research, 69(20), 8150–8156. doi:10.1158/0008-5472.CAN-09-1135

Eto, T., Miyake, K., Nosho, K., Ohmuraya, M., Imamura, Y., Arima, K., … Ishimoto, T. (2018). Impact of loss-of-function mutations at the RNF43 locus on colorectal cancer development and progression. J Pathol, 245(4), 445–455. doi:10.1002/path.5098

Furukawa, T., Kuboki, Y., Tanji, E., Yoshida, S., Hatori, T., Yamamoto, M., … Shiratori, K. (2011). Whole-exome sequencing uncovers frequent GNAS mutations in intraductal papillary mucinous neoplasms of the pancreas. Scientific reports, 1, 161. doi:10.1038/srep00161

Hua, X., Zhao, W., Pesatori, A. C., Consonni, D., Caporaso, N. E., Zhang, T., … Landi, M. T. (2020). Genetic and epigenetic intratumor heterogeneity impacts prognosis of lung adenocarcinoma. Nature communications, 11(1), 2459. doi:10.1038/s41467-020-16295-5

Iacobuzio-Donahue, C. A., Wilentz, R. E., Argani, P., Yeo, C. J., Cameron, J. L., Kern, S. E., & Hruban, R. H. (2000). Dpc4 protein in mucinous cystic neoplasms of the pancreas: frequent loss of expression in invasive carcinomas suggests a role in genetic progression. Am J Surg Pathol, 24(11), 1544–1548.

Ishidate, T., Matsumine, A., Toyoshima, K., & Akiyama, T. (2000). The APC-hDLG complex negatively regulates cell cycle progression from the G0/G1 to S phase. Oncogene, 19(3), 365–372. doi:10.1038/sj.onc.1203309

Jackson, E. L., Willis, N., Mercer, K., Bronson, R. T., Crowley, D., Montoya, R., … Tuveson, D. A. (2001). Analysis of lung tumor initiation and progression using conditional expression of oncogenic K-ras. Genes & development, 15(24), 3243–3248. doi:10.1101/gad.943001

Jamal-Hanjani, M., Quezada, S. A., Larkin, J., & Swanton, C. (2015). Translational implications of tumor heterogeneity. Clinical cancer research: an official journal of the American Association for Cancer Research, 21(6), 1258–1266. doi:10.1158/1078-0432.CCR-14-1429

Kang, Y., Zhang, R., Suzuki, R., Li, S. Q., Roife, D., Truty, M. J., … Fleming, J. B. (2015). Two-dimensional culture of human pancreatic adenocarcinoma cells results in an irreversible transition from epithelial to mesenchymal phenotype. Lab Invest, 95(2), 207–222. doi:10.1038/labinvest.2014.143

Kim, M. P., Evans, D. B., Wang, H., Abbruzzese, J. L., Fleming, J. B., & Gallick, G. E. (2009). Generation of orthotopic and heterotopic human pancreatic cancer xenografts in immunodeficient mice. Nature protocols, 4(11), 1670–1680. doi:10.1038/nprot.2009.171

Kuleshov, M. V., Jones, M. R., Rouillard, A. D., Fernandez, N. F., Duan, Q., Wang, Z., … Ma’ayan, A. (2016). Enrichr: a comprehensive gene set enrichment analysis web server 2016 update. Nucleic acids research. doi:10.1093/nar/gkw377

Laprise, P., Viel, A., & Rivard, N. (2004). Human homolog of disc-large is required for adherens junction assembly and differentiation of human intestinal epithelial cells. The Journal of biological chemistry, 279(11), 10157–10166. doi:10.1074/jbc.M309843200

Law, C. W., Chen, Y., Shi, W., & Smyth, G. K. (2014). voom: Precision weights unlock linear model analysis tools for RNA-seq read counts. Genome Biol, 15(2), R29. doi:10.1186/gb-2014-15-2-r29

Liao, Y., Smyth, G. K., & Shi, W. (2014). featureCounts: an efficient general purpose program for assigning sequence reads to genomic features. Bioinformatics, 30(7), 923–930. doi:10.1093/bioinformatics/btt656

Mann, K. M., Newberg, J. Y., Black, M. A., Jones, D. J., Amaya-Manzanares, F., Guzman-Rojas, L., … Mann, M. B. (2016). Analyzing tumor heterogeneity and driver genes in single myeloid leukemia cells with SBCapSeq. Nat Biotechnol, 34(9), 962–972. doi:10.1038/nbt.3637

Mann, K. M., Ward, J. M., Yew, C. C., Kovochich, A., Dawson, D. W., Black, M. A., … Copeland, N. G. (2012). Sleeping Beauty mutagenesis reveals cooperating mutations and pathways in pancreatic adenocarcinoma. Proceedings of the National Academy of Sciences of the United States of America, 109(16), 5934–5941. doi:10.1073/pnas.1202490109

Matsumine, A., Ogai, A., Senda, T., Okumura, N., Satoh, K., Baeg, G. H., … Akiyama, T. (1996). Binding of APC to the human homolog of the Drosophila discs large tumor suppressor protein. Science, 272(5264), 1020–1023. doi:10.1126/science.272.5264.1020

McDonald, K. A., Kawaguchi, T., Qi, Q., Peng, X., Asaoka, M., Young, J., … Takabe, K. (2019). Tumor Heterogeneity Correlates with Less Immune Response and Worse Survival in Breast Cancer Patients. Annals of surgical oncology, 26(7), 2191–2199. doi:10.1245/s10434-019-07338-3

Moffitt, R. A., Marayati, R., Flate, E. L., Volmar, K. E., Loeza, S. G., Hoadley, K. A., … Yeh, J. J. (2015). Virtual microdissection identifies distinct tumor-and stroma-specific subtypes of pancreatic ductal adenocarcinoma. Nature genetics, 47(10), 1168–1178. doi:10.1038/ng.3398

Mroz, E. A., Tward, A. D., Pickering, C. R., Myers, J. N., Ferris, R. L., & Rocco, J. W. (2013). High intratumor genetic heterogeneity is related to worse outcome in patients with head and neck squamous cell carcinoma. Cancer, 119(16), 3034–3042. doi:10.1002/cncr.28150

Network, C. G. A. R. (2017). Integrated Genomic Characterization of Pancreatic Ductal Adenocarcinoma. Cancer cell, 32(2), 185–203 e113. doi:10.1016/j.ccell.2017.07.007

Neumeyer, V., Grandl, M., Dietl, A., Brutau-Abia, A., Allgauer, M., Kalali, B., … Gerhard, M. (2019). Loss of endogenous RNF43 function enhances proliferation and tumour growth of intestinal and gastric cells. Carcinogenesis, 40(4), 551–559. doi:10.1093/carcin/bgy152

Newberg, J. Y., Black, M. A., Jenkins, N. A., Copeland, N. G., Mann, K. M., & Mann, M. B. (2018). SB Driver Analysis: a Sleeping Beauty cancer driver analysis framework for identifying and prioritizing experimentally actionable oncogenes and tumor suppressors. Nucleic acids research, 46(16), e94. doi:10.1093/nar/gky450

Newberg, J. Y., Mann, K. M., Mann, M. B., Jenkins, N. A., & Copeland, N. G. (2018). SBCDDB: Sleeping Beauty Cancer Driver Database for gene discovery in mouse models of human cancers. Nucleic acids research, 46(D1), D1011–D1017. doi:10.1093/nar/gkx956

Ngan, E., Stoletov, K., Smith, H. W., Common, J., Muller, W. J., Lewis, J. D., & Siegel, P. M. (2017). LPP is a Src substrate required for invadopodia formation and efficient breast cancer lung metastasis. Nat Commun, 8, 15059. doi:10.1038/ncomms15059

Niknafs, N., Zhong, Y., Moral, J. A., Zhang, L., Shao, M. X., Lo, A., … Karchin, R. (2019). Characterization of genetic subclonal evolution in pancreatic cancer mouse models. Nature communications, 10(1), 5435. doi:10.1038/s41467-019-13100-w

Novellino, L., De Filippo, A., Deho, P., Perrone, F., Pilotti, S., Parmiani, G., & Castelli, C. (2008). PTPRK negatively regulates transcriptional activity of wild type and mutated oncogenic beta-catenin and affects membrane distribution of beta-catenin/E-cadherin complexes in cancer cells. Cell Signal, 20(5), 872–883. doi:10.1016/j.cellsig.2007.12.024

Perez-Mancera, P. A., Rust, A. G., van der Weyden, L., Kristiansen, G., Li, A., Sarver, A. L., … Tuveson, D. A. (2012). The deubiquitinase USP9X suppresses pancreatic ductal adenocarcinoma. Nature, 486(7402), 266–270. doi:10.1038/nature11114

Qu, C., He, Lu, X., Dong, L., Zhu, Y., Zhao, Q., … Zhang, Z. (2016). Salt-inducible Kinase (SIK1) regulates HCC progression and WNT/beta-catenin activation. J Hepatol, 64(5), 1076–1089. doi:10.1016/j.jhep.2016.01.005

Rad, R., Rad, L., Wang, W., Strong, A., Ponstingl, H., Bronner, I. F., … Bradley, A. (2015). A conditional piggyBac transposition system for genetic screening in mice identifies oncogenic networks in pancreatic cancer. Nature genetics, 47(1), 47–56. doi:10.1038/ng.3164

Rangel, R., Lee, S. C., Hon-Kim Ban, K., Guzman-Rojas, L., Mann, M. B., Newberg, J. Y., … Copeland, N. G. (2016). Transposon mutagenesis identifies genes that cooperate with mutant Pten in breast cancer progression. Proceedings of the National Academy of Sciences of the United States of America, 113(48), E7749–E7758. doi:10.1073/pnas.1613859113

Reichert, M., Bakir, B., Moreira, L., Pitarresi, J. R., Feldmann, K., Simon, L., … Rustgi, A. K. (2018). Regulation of Epithelial Plasticity Determines Metastatic Organotropism in Pancreatic Cancer. Dev Cell, 45(6), 696–711 e698. doi:10.1016/j.devcel.2018.05.025

Reuben, A., Spencer, C. N., Prieto, P. A., Gopalakrishnan, V., Reddy, S. M., Miller, J. P., … Wargo, J. A. (2017). Genomic and immune heterogeneity are associated with differential responses to therapy in melanoma. NPJ Genom Med, 2. doi:10.1038/s41525-017-0013-8

Sakamoto, H., Kuboki, Y., Hatori, T., Yamamoto, M., Sugiyama, M., Shibata, N., … Furukawa, T. (2015). Clinicopathological significance of somatic RNF43 mutation and aberrant expression of ring finger protein 43 in intraductal papillary mucinous neoplasms of the pancreas. Mod Pathol, 28(2), 261–267. doi:10.1038/modpathol.2014.98

Setzer, S. V., Calkins, C. C., Garner, J., Summers, S., Green, K. J., & Kowalczyk, A. P. (2004). Comparative analysis of armadillo family proteins in the regulation of a431 epithelial cell junction assembly, adhesion and migration. J Invest Dermatol, 123(3), 426–433. doi:10.1111/j.0022-202X.2004.23319.x

Siegel, R. L., Miller, K. D., & Jemal, A. (2019). Cancer statistics, 2019. CA: a cancer journal for clinicians, 69(1), 7–34. doi:10.3322/caac.21551

Sveen, A., Loes, I. M., Alagaratnam, S., Nilsen, G., Holand, M., Lingjaerde, O. C., … Lothe, R. A. (2016). Intra-patient Inter-metastatic Genetic Heterogeneity in Colorectal Cancer as a Key Determinant of Survival after Curative Liver Resection. PLoS Genet, 12(7), e1006225. doi:10.1371/journal.pgen.1006225

Szklarczyk, D., Gable, A. L., Lyon, D., Junge, A., Wyder, S., Huerta-Cepas, J., … Mering, C. V. (2019). STRING v11: protein-protein association networks with increased coverage, supporting functional discovery in genome-wide experimental datasets. Nucleic acids research, 47(D1), D607–D613. doi:10.1093/nar/gky1131

Takeda, H., Rust, A. G., Ward, J. M., Yew, C. C., Jenkins, N. A., & Copeland, N. G. (2016). Sleeping Beauty transposon mutagenesis identifies genes that cooperate with mutant Smad4 in gastric cancer development. Proceedings of the National Academy of Sciences of the United States of America. doi:10.1073/pnas.1603223113

The Gene Ontology, C. (2019). The Gene Ontology Resource: 20 years and still GOing strong. Nucleic acids research, 47(D1), D330–D338. doi:10.1093/nar/gky1055

Waddell, N., Pajic, M., Patch, A. M., Chang, D. K., Kassahn, K. S., Bailey, P., … Grimmond, S. M. (2015). Whole genomes redefine the mutational landscape of pancreatic cancer. Nature, 518(7540), 495–501. doi:10.1038/nature14169

Wang, X., Spandidos, A., Wang, H., & Seed, B. (2012). PrimerBank: a PCR primer database for quantitative gene expression analysis, 2012 update. Nucleic acids research, 40(Database issue), D1144–1149. doi:10.1093/nar/gkr1013

Wang, Z., Hausmann, S., Lyu, R., Li, T. M., Lofgren, S. M., Flores, N. M., … Mazur, P. K. (2020). SETD5-Coordinated Chromatin Reprogramming Regulates Adaptive Resistance to Targeted Pancreatic Cancer Therapy. Cancer cell, 37(6), 834–849 e813. doi:10.1016/j.ccell.2020.04.014

Welsh, E. A., Eschrich, S. A., Berglund, A. E., & Fenstermacher, D. A. (2013). Iterative rank-order normalization of gene expression microarray data. BMC bioinformatics, 14, 153. doi:10.1186/1471-2105-14-153

Witkiewicz, A. K., McMillan, E. A., Balaji, U., Baek, G., Lin, W. C., Mansour, J., … Knudsen, E. S. (2015). Whole-exome sequencing of pancreatic cancer defines genetic diversity and therapeutic targets. Nature communications, 6, 6744. doi:10.1038/ncomms7744

Wood, L. D., Parsons, D. W., Jones, S., Lin, J., Sjoblom, T., Leary, R. J., … Vogelstein, B. (2007). The genomic landscapes of human breast and colorectal cancers. Science, 318(5853), 1108–1113. doi:10.1126/science.1145720

Yachida, S., Jones, S., Bozic, I., Antal, T., Leary, R., Fu, B., … Iacobuzio-Donahue, C. A. (2010). Distant metastasis occurs late during the genetic evolution of pancreatic cancer. Nature, 467(7319), 1114–1117. doi:10.1038/nature09515

Yaeger, R., Chatila, W. K., Lipsyc, M. D., Hechtman, J. F., Cercek, A., Sanchez-Vega, F., … Schultz, N. (2018). Clinical Sequencing Defines the Genomic Landscape of Metastatic Colorectal Cancer. Cancer cell, 33(1), 125–136 e123. doi:10.1016/j.ccell.2017.12.004

Young, M. D., Wakefield, M. J., Smyth, G. K., & Oshlack, A. (2010). Gene ontology analysis for RNA-seq: accounting for selection bias. Genome Biol, 11(2), R14. doi:10.1186/gb-2010-11-2-r14

Zhang, J., Bajari, R., Andric, D., Gerthoffert, F., Lepsa, A., Nahal-Bose, H., … Ferretti, V. (2019). The International Cancer Genome Consortium Data Portal. Nat Biotechnol, 37(4), 367–369. doi:10.1038/s41587-019-0055-9

Zhang, J., Tsoi, H., Li, X., Wang, H., Gao, J., Wang, K., … Yu, J. (2016). Carbonic anhydrase IV inhibits colon cancer development by inhibiting the Wnt signalling pathway through targeting the WTAP-WT1-TBL1 axis. Gut, 65(9), 1482–1493. doi:10.1136/gutjnl-2014-308614

Zhao, B., Hemann, M. T., & Lauffenburger, D. A. (2014). Intratumor heterogeneity alters most effective drugs in designed combinations. Proceedings of the National Academy of Sciences of the United States of America, 111(29), 10773–10778. doi:10.1073/pnas.1323934111

